# Why do young readers vary in reading fluency? The impact of word length and frequency in French 6th graders

**DOI:** 10.1101/2023.01.30.526188

**Authors:** Marie Lubineau, Cassandra Potier Watkins, Hervé Glasel, Stanislas Dehaene

## Abstract

**Purpose:** Which processes induce variations in reading speed in young readers with the same amount of education, but different levels of reading fluency? Here, we tested a prediction of the dual-route model: as fluency increases, these variations may reflect a decreasing reliance on decoding and an increasing reliance on the lexical route.

**Method:** 1,500 French 6^th^ graders passed a one-minute speeded reading-aloud task evaluating fluency, and a 10-minute computerized lexical decision task evaluating the impact of word length, word frequency and pseudoword type.

**Results:** As predicted, the word length effect varied dramatically with reading fluency, with the least fluent group showing a length effect even for frequent words. The frequency effect also varied, but solely in proportion to overall slowness, suggesting that frequency affects the decision stage in all readers, while length impacts poor readers disproportionately. Response times and errors were also affected by pseudoword type (e.g. letter substitutions or transpositions), but these effects did not vary much with fluency. Overall, lexical decision variables were excellent predictors of reading fluency (r=0.62).

**Conclusion:** Our results call attention to middle-school reading difficulties and encourage the use of lexical decision as a test of students’ mental lexicon and the automatization of reading.

## Introduction

Why does reading performance vary so much across students from the same grade? And could a brief in-school test shed light on the origins of their reading difficulties, thus helping teachers provide them with appropriate feedback? In France, where the present study took place, as in many other countries, many students enter middle school without the skills needed for independent reading (Andreu et al., 2021, 2022). It is in this context that the French government initiated a series of National Evaluations at the start of the 6^th^ grade (first year of middle school in France), with the goal to provide teachers with individualized information about their students’ reading difficulties. The National Evaluations are computer-based, except for a one-minute read-aloud text reading fluency test. Such a fluency test assesses fast and accurate text reading, necessitating appropriate prosody or stress (Hudson et al., 2005), which is considered as a hallmark of reading automaticity (Breznitz, 1987; LaBerge & Samuels, 1974; National Reading Panel, 2000; Pinnell et al., 1995) and strongly correlates with reading comprehension, the goal of reading (Fuchs et al., 2001). One criticism of fluency assessment, however, is that it calls for the mapping of orthographic input to phonological output, which is unnecessary for silent reading. The Lexical Decision (LD), a classic psycholinguistics paradigm requiring participants to classify visually presented stimuli as words or pseudowords, has emerged as an alternative assessment (Gijsel et al., 2004; van Bon et al., 2004; Yeatman et al., 2021) possibly more diagnostic of the orthographic-to-lexical route involved in silent reading comprehension (Holmes, 2009; Martens & de Jong, 2006; Perfetti, 2007). Here, we investigate whether a fast computerized LD task may shed light on the origins of inter-individual variability in reading fluency.

The LD task has been widely used in cognitive science and provides well-replicated measures of key components of the visual word recognition process, which are captured by the Dual Route Model of reading (Coltheart et al., 2001; Fiebach et al., 2002). This model describes the reading process as the outcome of two competing pathways. On the one hand, the lexical pathway recognizes words already stored in the mental lexicon. This lexicon is affected by word frequency, such that most frequent words are quicker to retrieve than less frequent ones. This pathway lends itself to fast fluent reading of known words. On the other hand, the sublexical pathway enables reading words that have not been stored in the mental lexicon, such as unknown words or pseudowords, via the application of grapheme-phoneme correspondence (GPC) rules. The necessity to decode graphemes into phonemes in order to access sound and thereby meaning (or rejection of meaning) causes response time (RT) to stimuli processed via the sublexical pathway to be strongly affected by a length effect (Acha & Perea, 2008; Bijeljac-Babic et al., 2004; Di Filippo et al., 2006; Martens & de Jong, 2006).

During LD, where the task is to make a judgement of an item’s lexicality, a lexicality effect is typically observed, with more accurate and faster responses to words than pseudowords or unknown words (Di Filippo et al., 2006). The superiority of the lexical pathway, however, is mitigated by word frequency -- actually a mixture of age of acquisition and frequency of use (Brysbaert et al., 2011; Grainger & Segui, 1990; Juhasz et al., 2019; Morrison & Ellis, 2000).

The variables that affect LD capture the changing weights of the two pathways during reading development and can therefore provide researchers and educators with valuable measures for reading assessment (Yeatman et al., 2021). Children, compared to adults, are affected by a more pronounced word length effect, as they rely more heavily on sublexical procedures when learning to read (Acha & Perea, 2008). As reading becomes efficient, the word length effect fades, indicating that the lexical pathway becomes fully operational (Weekes, 1997; Zoccolotti et al., 2005). The lexicality effect generally appears for high frequency words by 3^rd^ grade (Araújo et al., 2014; Di Filippo et al., 2006; Juphard et al., 2004; Sela et al., 2014). However, this maturing of faster lexical procedures can take from 1 to 2 years, depending on the transparency of the orthography in a given language (Schmalz et al., 2013). In French, a language with an opaque orthography that comprises many graphemes with different possible phonemic realizations, many rules must be memorized in order to decode all words correctly. It takes French children almost a year of reading acquisition for the frequency effect to appear (Sprenger-Charolles et al., 1998). The word length effect diminishes between 3^rd^ and 5^th^ grade, indicating a slow transition from serial grapheme–phoneme mapping to a greater reliance on lexical knowledge (Bijeljac-Babic et al., 2004). This lack of orthographic transparency modulates the speed with which children learn to read, as well as the types of deficits that manifest in dyslexia (Ziegler et al., 2010).

The decrease in the length effect is concomitant with the appearance of a frequency effect (Faust, Balota, Spieler, Ferraro, et al., 1999). Word reading performance improves for longer words, and eventually ceases to increase with length over a broad range of ∼3-8 letters (New et al., 2006), while RT and accuracy become increasingly affected by frequency (Brysbaert et al., 2011; Burani et al., 2002; Grainger & Segui, 1990; Ratcliff et al., 2004). At this stage, reading is characterized by a linear relationship between the logarithm of the frequency and RT (Norris, 2006).

Studies in dyslexic readers further supports LD as a useful task to measure the developmental trajectory of reading. When compared to chronological age controls, dyslexic readers show slower RT and larger effects of word length and lexicality, similar to younger children matched in terms of raw reading scores (Castles, 2006; Martens & de Jong, 2006; Zoccolotti et al., 2005). However, these older dyslexic students are closer to their age matched controls in terms of frequency effects (Martens & de Jong, 2006).

### The current project

The evidence that LD can capture variations in the pathways for visual word recognition is clear, but the major issue remains whether LD can provide an informative assessment of reading skills. Previous studies have had mixed results (Gijsel et al., 2004; van Bon et al., 2004; Yeatman et al., 2021). In second and third graders, LD is highly correlated with other standardized oral reading outcomes, with high test-retest reliability (van Bon et al., 2004). There is a decline, however, in the correlation of LD with standardized oral reading in readers from 3^rd^ to 6^th^ grade (Gijsel et al., 2004; van Bon et al., 2004). In adults, LD does not appear to predict reading comprehension (Katz et al., 2012). However, in the Katz and collaborators research, all the LD words were frequent and adult accuracy was at ceiling, meaning that there was possibly not enough variability to get an accurate picture of ability. Which LD outcomes should be measured is also unclear. Compared to adults, accuracy appears to be a better predictor of oral word reading than RT for children, with pseudoword responses more informative than words (Yeatman et al., 2021).

The primary goal of the current study was to extend our knowledge of LD as a measure of the underlying processes in silent reading, with a particular focus on lexicality, length, and frequency effects. Prior research on LD as a tool for reading assessment has used it to correlate it with other reading tests in adults (Katz et al., 2012; Yeatman et al., 2021); to probe grade-level differences in children (Gijsel et al., 2004; van Bon et al., 2004; Yeatman et al., 2021); or to compare, within a given grade, normal readers with dyslexic students (Araújo et al., 2014; Di Filippo et al., 2006; Juphard et al., 2004; Zoccolotti et al., 2005). Novel to our approach, we examine within-grade variability on a LD task. We obtained LD data from a large sample of students in 6^th^ grade (N= 1,500, mean age of 11y.o.), which is the first year of middle school in France, a turning point given the increasing importance of silent reading skills and decreasing teacher-led explicit reading instruction. We subdivided our participants based on groups of reading ability from their national fluency scores, then investigated how these groups perform on a LD task. We provide a detailed account of group lexical and sublexical reading procedures through the prism of accuracy and RT. Given that students in this grade have had five years of reading exposure, we hypothesized that, ideally, all students should show no length effect for frequent words, and that length effects would persist for less frequent and rare words, predominantly for poor readers.

Another goal of our work was to better understand how students at different levels of reading ability process pseudowords. To this aim, we designed different types of word-derived pseudowords, also called “traps” because they must be rejected, in spite of their often close similarity to words. Much prior research has demonstrated an effect of the orthographic similarity of pseudowords to words (Davis & Bowers, 2006; Ferrand & Grainger, 1993; Grainger & Segui, 1990). This similarity is classically measured using Coltheart’s N (Coltheart et al., 1977), which considers as “neighbors” two strings of the same length that differ by only one letter. Here, however, we explored a greater range of pseudoword types, based on the presence of transposed or mirror letters, as well as misspelled words.

Developing as well as skilled readers often make errors that involve transpositions of internal letters (for example “from” and “form”) (Paterson et al., 2015). Children are also more likely to miscategorize transposed-letter pseudowords in LD tasks, an effect that first increased with reading acquisition and then decreases to its minimum for skilled readers (Grainger et al., 2012). We expected our participants to be slower and less accurate when processing letter-transposition pseudowords than their double-substitution controls, an effect that would be highest for most fluent readers. Regarding mirror generalization, it is an early predisposition of the pre-reader’s brain that must be inhibited or superseded when learning to read (Dehaene, 2009; Dehaene et al., 2015; Kolinsky et al., 2011; Pegado et al., 2011, 2014). Indeed, identifying that two mirror letters are different is harder than differentiating between two non-mirror letters (Ahr et al., 2016). In addition, the sounds /p/-/b/ and /b/-/d/ are very close phonologically. Thus, we expected that processing pseudowords containing mirror substitutions, such as ‘dateau’ instead of the French word ‘bateau’, would require a greater effort than processing pseudowords arising from an equivalent, non-mirror letter substitution (e.g. ‘fateau’; English equivalents would be ‘dalance’ [derived from ‘balance’] versus ‘falance’). As children with reading deficits often confuse letter-sound rules (Rack et al., 1992), we also introduced misspelled words that would sound like a word if the wrong GPC rule was applied (pseudohomophones). Prior research using lexical decision showed that students exhibited major difficulties distinguishing between words and pseudohomophones (Bergmann & Wimmer, 2008) which decreased in the course of reading development (Grainger et al., 2012). Thus, we expected higher RTs and error rates for orthographic traps than for control word approximations, and a reduction of this effect as fluency increased. Errors on pseudohomophones may be an outcome of poor reading experience exasperated by difficulties in learning French, a language with and opaque orthography. We hypothesized that all readers would be slower to classify pseudowords based on their distance to a real word, but that poor readers would be further penalized by pseudohomophones.

Finally, our last goal was to investigate the predictive value of the LD task on fluency. LD was shown to be highly correlated with a standardized measure of oral reading for children and adults (Yeatman et al., 2021, Gijsel et al., 2004; van Bon et al., 2004)). Here, we investigated how LD accuracy and RT varied in relation to text reading fluency. By correlating these two tasks, we aimed to further our understanding of how elementary measures of single-word phonological and orthographic processing, acquired during LD, relate to fluent text processing.

## Methods

### Participants

In the context of the French national evaluations given to all students in 6th grade (the first year of middle school), we deployed our LD task in a panel of students. Test administration and data collection were performed by the ministerial service in charge of education statistics, the Direction of Evaluation, Prospective and Performance (DEPP, www.education.gouv.fr/direction-de-l-evaluation-de-la-prospective-et-de-la-performance-depp-12389). 3,614 students, selected by the DEPP as representative population of France, were included in the panel. Due to a software error (leading to missing data and negative RTs in some students) only 1,500 participants with a complete dataset were retained for our analysis (806 girls and 694 boys, mean age = 11.0 y.o.). We received from the DEPP anonymized data from a standardized fluency test and our LD task.

### Reading fluency test

This portion of the national evaluation was administered individually by the teacher. In a quiet room, the student was instructed to read aloud a fixed, grade-appropriate text for one minute, as accurately as possible in normal reading speed. Teachers reported the number of words correctly read in one minute by each child.

### Lexical Decision Task

The LD task was included in the computerized national assessment. Students worked individually with headphones in a group setting in the school’s computer lab. The task started with written and oral instructions: “For this exercise, decide as fast as possible if what is written on the screen is a real word or a trap. Press M for a word and Q for a trap.” This was followed by a video demonstration for each item category (word or trap). Students clicked a button when they were ready to start. Each item remained in the middle of the screen until the student responded by pressing ‘Q’ or ‘M’ or was skipped after a 5000ms time limit. Audio-visual feedback was provided. Positive audio feedback increased in tone with consecutive correct responses to encourage pursuit of winnings streaks. We collected measures of accuracy and response time (RT in ms). There were twelve different lexical decision modules, each composed of 120 stimuli, 60 words and 60 pseudowords (see below). Students were randomly assigned to a module. The whole task was administered in one block, and stimulus order was randomized within each student.

### Word stimuli

We first extracted all mono-lemmatic and mono-morphemic words from the Lexique 3.83 database (New et al., 2004) with a length of four to eight letters and a frequency higher than three per million. We manually excluded all potentially offending, inappropriate or foreign words, thus resulting in a stimulus set of 3,656 words. These items were then separated into four different frequency bands: very frequent, frequent, rare and very rare (see table 1 for details). 12 modules were designed using this database. For each module, we randomly selected 3 words from each frequency category and each length, resulting in a factorial design with length (5 levels, 4-8 letters) and frequency (4 levels) as factors, and a total of 60 words per module.

**Table 1.**
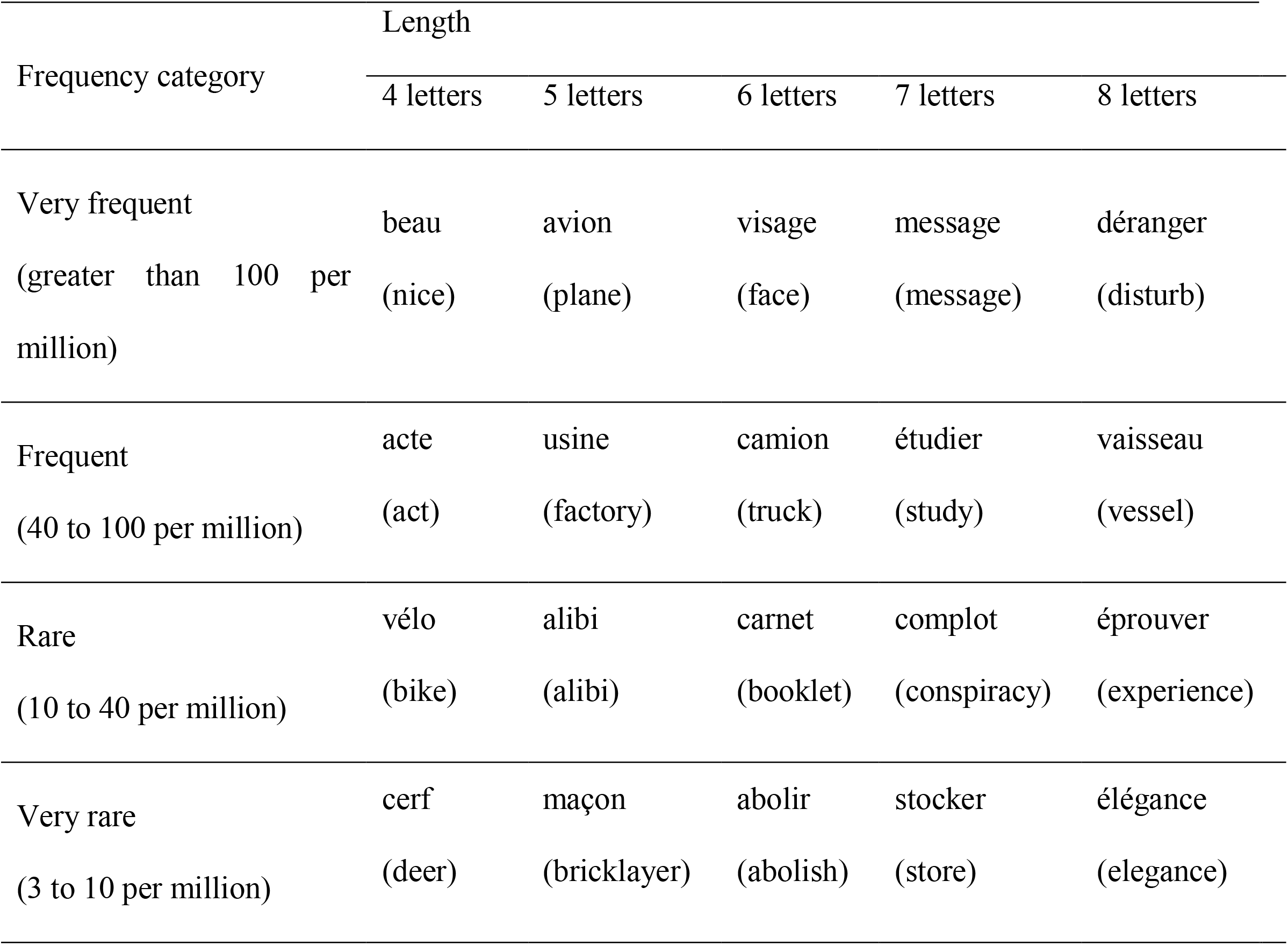
Characteristics of words stimuli and examples.

### Pseudoword stimuli

Starting from words from the “very frequent” category, we built six categories of pseudowords *“traps”*, for a total pool of 1,196 items. For each module, we randomly selected two pseudowords for each type and each length, thus resulting in a factorial design with pseudoword category (6 levels, see below) and length (5 levels, 4-8 letters) as factors, and 60 pseudowords per module. We describe each of the pseudoword categories below. Examples are presented in Table 2.

**Table 2.** Characteristics of pseudowords stimuli and examples.

| Pseudoword traps | Length |  |  |  |  |
| --- | --- | --- | --- | --- | --- |
|  | 4 letters | 5 letters | 6 letters | 7 letters | 8 letters |
| Orthographic traps | bage | plase | inciet | ésaiyer | difisile |
|  | (bague = ring) | (place = place) | (inquiet = worried) | (essayer = try) | (difficile = hard) |
| Word approximations | atio | ouvoi | jamure | répoure | voicider |
| Transpositions | ceil | juene | geurre | pafrois | regadrer |
|  | (ciel = sky) | (jeune = young) | (guerre = war) | (parfois = sometimes) | (regarder = look) |
| Double substitutions | cuol | jaine | gairre | pansois | reganger |
|  | (ciel = sky) | (jeune = young) | (guerre = war) | (parfois = sometimes) | (regarder = look) |
| Mirror substitutions | aibe | qièce | dateau | musipue | qrochain |
|  | (aide = help) | (pièce = room) | (bateau = boat) | (musique = music) | (prochain = next) |
| Single substitutions | aite | gièce | fateau | musijue | grochain |
|  | (aide = help) | (pièce = room) | (bateau = boat) | (musique = music) | (prochain = next) |

#### Orthographic traps

These were misspelled words that could, by an erroneous grapheme-phoneme correspondence, sound as a real word. These pseudowords were manually built by selecting all words in our pool with a given rule-based grapheme-phoneme correspondence, and then over-regularizing it. We focused on the letters s, c, and g, whose pronunciation varies with context in French. The letter ‘s’ most frequently sounds as /s/ (e.g. in “sale”) except when a single ‘s’ is sandwiched between vowels (e.g. in “base”), causing it to sound /z/. Similarly, letters ‘c’ and ‘g’ respectively sound as /k/ and /g/ when followed by the vowels ‘a’, ‘o’ and ‘u’, but as /s/ and /Z/ when followed by the letters ‘e’ and ‘i’. As a result of those rules, the pseudoword ‘ausi’ should be read /ozi/, and the pseudoword ‘bage’ should be read /baZ/. A reader for whom these rules have not been consolidated might read these pseudowords as the words /osi/ (also) and /bag/ (ring)

#### Word approximations

Those were pseudowords entirely made of frequent French trigrams. To build them, we calculated the frequencies of all legal letter trigrams in our word pool (made of 26 letters plus an initial/final symbol). Only trigrams with frequency greater than 1/10,000 were retained. Pseudowords of 4 to 8 letters were then built solely from those frequent trigrams. We implemented a Markov process that (1) draws an initial trigram at random, in proportion to its frequency; (2) uses the last two letters to continue with the next trigram, again drawing randomly based on frequency, and so on. For example, the 4-letter pseudoword “arie” could be built using the frequent trigrams ‘#ar’, ‘ari’, ‘rie’, and ‘ie#’. Actual words were excluded by software and human inspection.

#### Letter transposition traps

Pseudo words of this category were constructed by inverting two adjacent internal letters of a word. The transposed bigram was composed of either two vowels or two consonants. Only pseudoword items whose bigrams exceeded a frequency of 1/10,0000 were kept. Bigram frequency was calculated using the same method described above for trigrams.

#### Two-letter substitution traps

This category, a control for transpositions, was built by substituting the same bigram in each transposed pseudoword with another random bigram with frequency higher than 1/10,000. Consonants were replaced by consonants, and vowels by vowels. Bigrams resulting from this substitution were all controlled to have a frequency higher than 1/10,000.

#### Mirror letter traps (mirroring of letters b d p q)

This category was generated by mirroring those letters in the following way: p→q ; q→p ; b→d ; d→b. Items were only kept if the transformation yielded a pseudoword.

#### Single-letter substitution traps (substitution of letters b d p q)

In this category, a control for the mirror substitution traps, letters b d p q were substituted with a non-mirror letter:

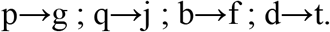

### Data Analyses

Trials with RT > 200ms were included into a linear mixed-effect model with fluency level (quintiles 1-5), lexicality (word, pseudoword), length (4-8 letters), word frequency (4 levels from very frequent to very rare), and pseudoword type (6 levels) as fixed effects, and subject and stimulus as random effects. We used the ‘mixed’ function from R’s afex package with the following formula:

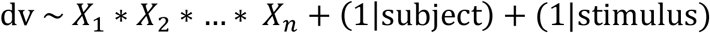

where the *X*_*i*_ are our fixed effects interacting together and *dv* is either accuracy or RT. For RT, we computed a classical mixed effect model (Baayen et al., 2008) while for accuracy we used a logistic mixed effect regression with the binomial link function (Jaeger, 2008). Significance was computed using the Satterthwaite’s degrees of freedom for RT and the Likelihood ratio test (LRT) for accuracy. All analyses used a significance threshold of α=0.05.

Follow-up analyses on significant interactions were done using the simple effect analysis where we split the data into subsets according to the modulating variable(s) and recomputed the model with only the remaining variable(s).

To investigate the predictive value of the LD on reading fluency, we used both simple and multiple linear regression. The predictive variables were the student’s median RT on correct answers across all trials, global error rate, slope of the length effect, slope of the frequency effect on words, and efficiency score (ratio of accuracy by mean RT).

## Results

Figure 1 shows the distribution of text reading fluency in our sample, in words/minutes. Since the primary goal of our project was to assess within-grade variability in the size of the lexicality, length and frequency effects in the LD task results, we first separated students into five groups based on their fluency score, each quintile group containing 20% of the children. Fluency quintile 1 refers to the best readers and 5 to the poorest readers.

**Figure 1.**
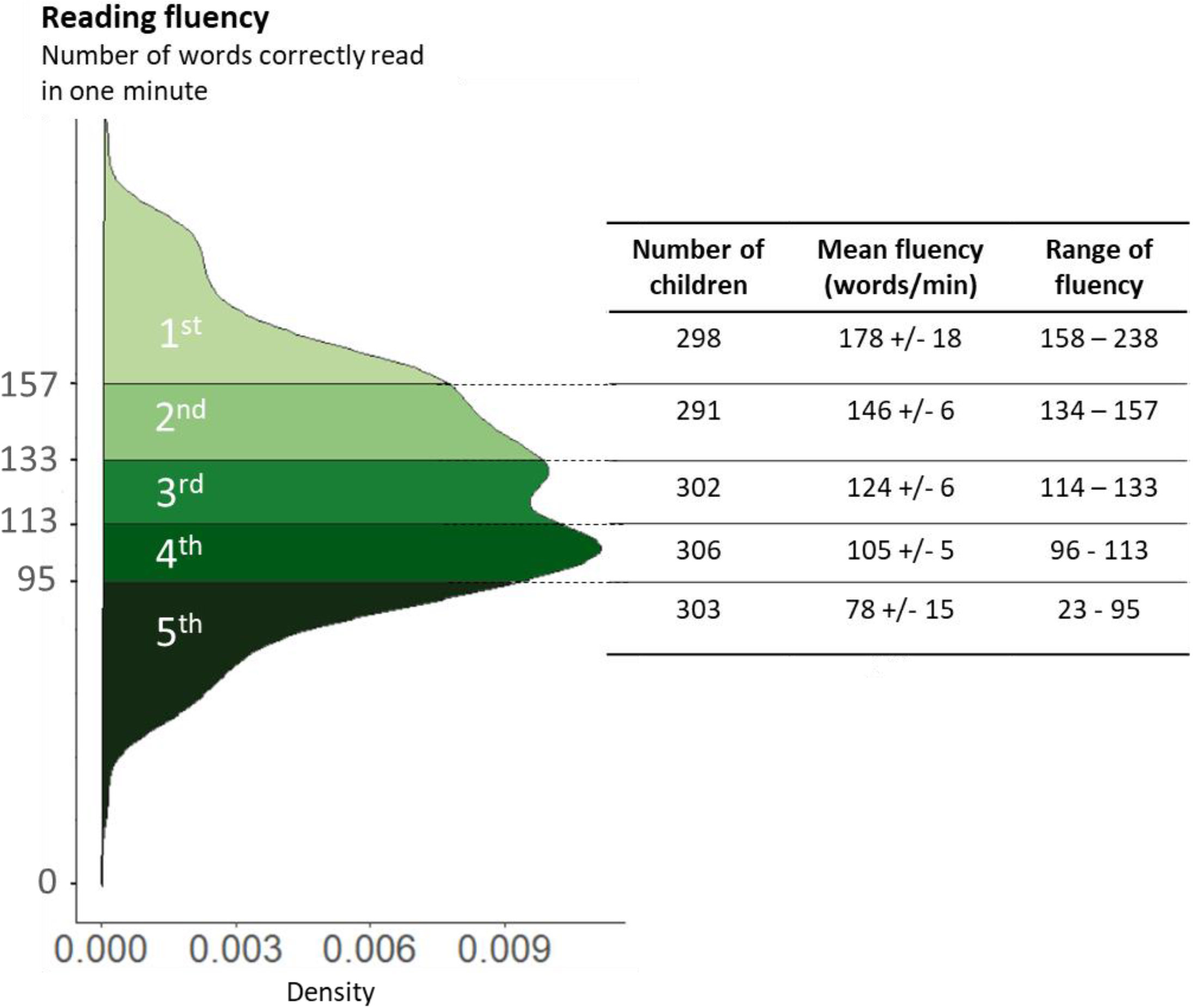
Distribution of participant’s reading fluency on the National Evaluation. Reading fluency was assessed by measuring the number of words correctly read in a text. In one minute. Color graduation from dark to light corresponds to fluency group levels, with light green corresponding to the most fluent students. Each quintile group represents 20% of the tested population.

### Length and lexicality effects

LD results are plotted in figure 2 as a function of lexical status and length. In agreement with the literature on LD RT, responses to words were faster than to pseudowords, F(1,1063.1)=212.82, p<.0001. There was a main effect of fluency quintile: the better the reader, the faster the LD RT, F(4,1478.2)=58.18, p<.0001. A main effect of length indicated that stimuli were read slower with increasing length, F(4,1062.7)=73.55, p<.0001. All three two-way interactions were significant: fluency x lexicality, F(4,135903.5)=16.49, p<.0001; fluency x length, F(4,135817.3)=50.13, p<.0001; lexicality x length, F(1,1062.8)=7.57, p=0.006. There was, however, no significant three-way interaction between fluency, lexicality, and length F(4,135823.9)=0.19, p=0.94.

**Figure 2.**
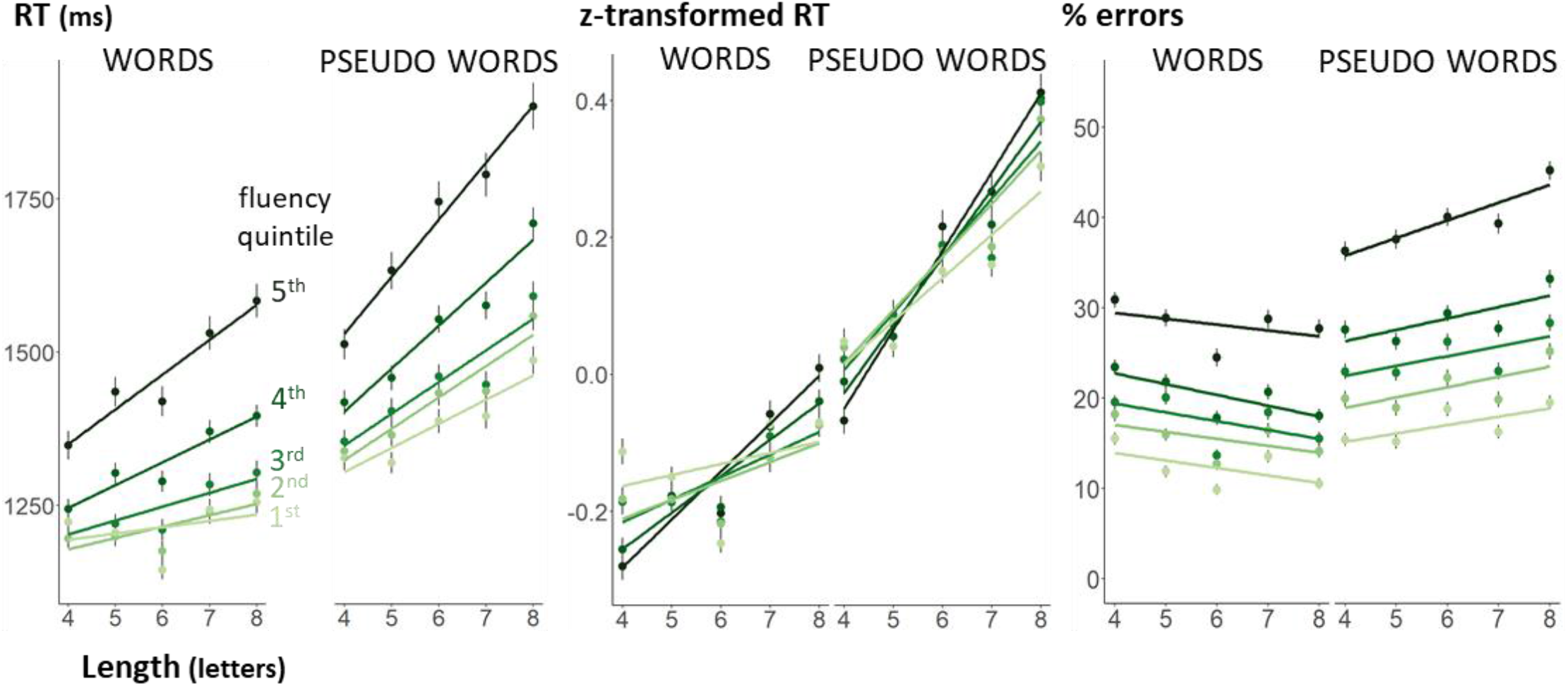
Length and lexicality effects on response times on correct answers (RT), Z-transformed RTs, and error rates. Each point represents the mean RT or error rate as function of word length and fluency. Error bars represent one standard error of the mean. The slopes are the linear regression associated with the points.

Simple effect analysis showed that the interaction between length and lexicality was significant within all fluency quintiles. Looking at each lexicality x fluency combination, we found that the length effect was significant in pseudowords no matter the fluency level, whereas for words, the length effect disappeared for the most fluent students, F(1,542.05)=3.00, p=0.084. When restricting the analysis to words only, there was a highly significant interaction of fluency by length, F(4,70591.24)=29.07, p<.001, confirming that the length effect vanished as fluency increased.

As expected, participants with lower fluency were also slower in the lexical decision task. Could this slowness alone explain why less fluent subjects showed a larger (absolute) effect of word length on RTs? Indeed, slower participants often shows larger RT effects than faster ones (Faust, Balota, Spieler, & Ferraro, 1999). If those effects occur at the decision stage, which is thought to involve a stochastic accumulation of evidence, then one would predict effect size to be proportional to the standard deviation of RTs across trials (Sigman & Dehaene, 2005). After verifying that our data fitted the conditions to apply Faust et al.’s (1999) rate and amount model (RAM), we turned our raw RT into z-scores by subtracting from each subject’s RT their overall mean and then dividing by their overall standard deviation (Zoccolotti et al., 2008). The results remain unchanged. Our mixed effect model on z-transformed RTs showed that all effects were significant except the three-way interaction, F(4,137191.6)=0.46, p=0.76. Crucially, within words, the fluency by length interaction remained significant, F(4,72020.47)=20.80, p<.0001, indicating that the reduction of the length effect as fluency increased went beyond what could be due solely to faster responses. Fluent readers are not just faster, but have a genuinely smaller length effect.

Results on error rates were close to those on RT. All subjects were more accurate in providing correct responses to words than pseudowords, χ²(1)=95.38, p<.001. Better readers were also more accurate in their decisions of lexicality, χ²(4)=593.03, p<.001. The main effect of length was not significant, χ²(1)=0.01, p=0.94, but a significant interaction was found between length and lexicality, χ²(1)=14.12, p<.001: greater length increased the likelihood of errors for pseudowords, and decreased it for words. We also found a significant interaction between fluency and lexicality, χ²(4)=24.34, p<.001 though not between fluency and length, χ²(4)=8.98, p=0.062, nor a three-way interaction, χ²(4)=1.09, p=0.90. We next investigated the length effect for each fluency x lexicality combination. All groups (except the 2^nd^ fluency quintile, χ²(1)=3.75, p=0.053) exhibited a positive length effect on pseudowords errors. For words, longer words tended to yield fewer errors, though this effect failed to reach significance for the 2^nd^ quintile, χ²(1)=3.49, p=0.062 and the 5^th^ quintile, χ²(1)=2.12, p=0.15.

### Error trials

All of the above analyses were performed on RTs to correct trials only, thus excluded many trials from the analysis, especially for the least fluent students. To investigate the processing of erroneous items, we applied the same mixed-effect analysis to error RTs (figure 3). We found a main effect of length F(1,1331.5)=28.29, p<.0001, as well as fluency F(4,1560.6)=3.83, p=0.0042. Looking at words and pseudowords separately, we observed that the length effect on words was the same for all fluency quintiles, as the interaction between both was not significant F(4,15791.17)=1.03, p=0.39. These results suggest that unknown words were treated serially as pseudowords, whatever the fluency level. For pseudowords, contrariwise, the interaction between length and fluency level was significant F(4,22253.14)=2.5, p=0.040, indicating a remarkable reduction and even disappearance of the length effect in the most fluent group. The latter finding suggests that these items were genuinely mistaken for words and processed non-serially through the lexical route.

**Figure 3.**
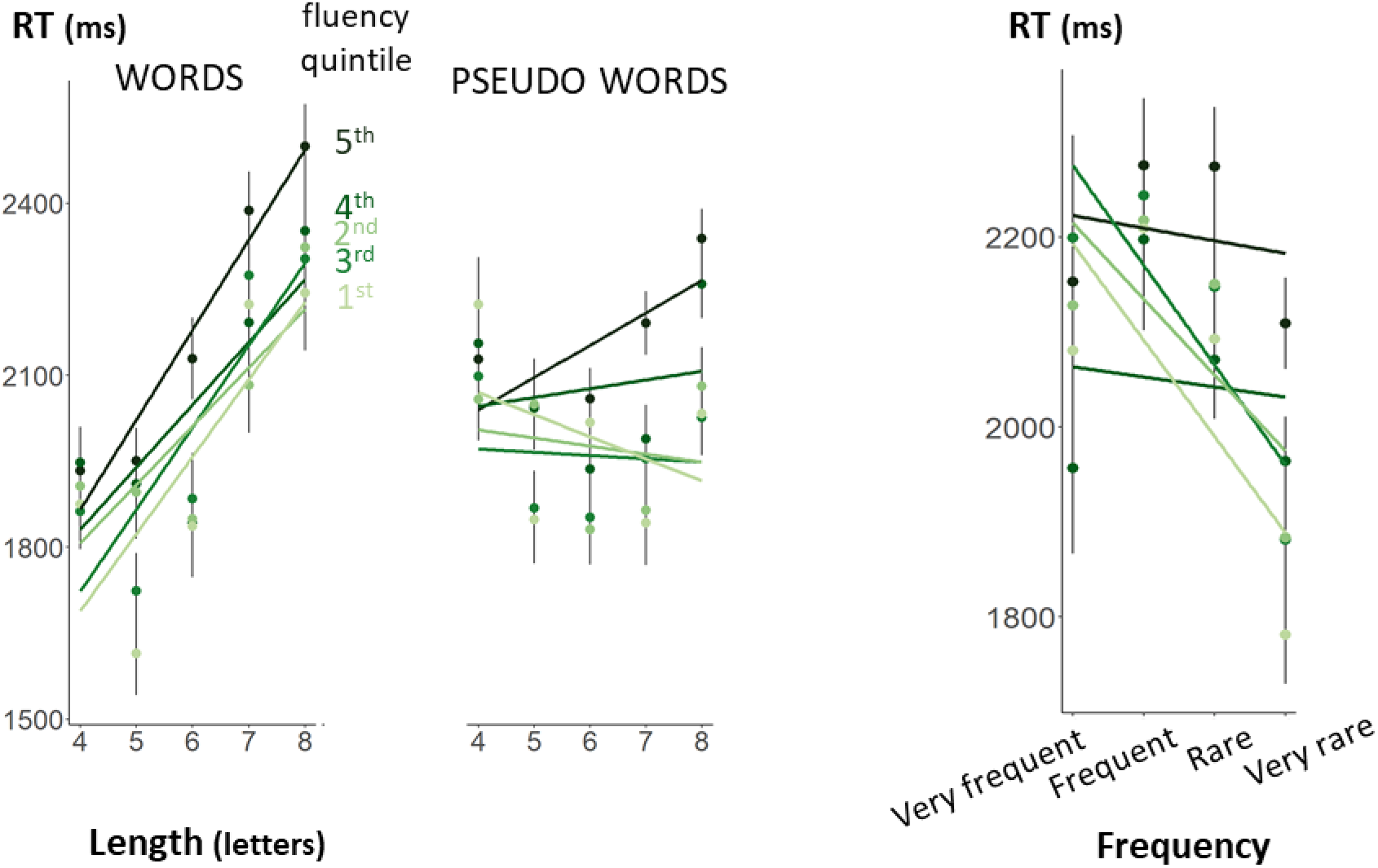
Length, lexicality and frequency effects on error trials. Each point represents the mean RT or error rate as a function of length and fluency, separately for words and pseudowords, solely on error trials. Error bars represent the standard error of the mean. The slopes are the linear regression associated with the points.

### Frequency effect

We next investigated the impact of word frequency and its variations with fluency. Correct RTs and error rates sorted by these four categories of frequency appear in figure 4. Both RT and accuracy were affected by a significant main effect of frequency (RT: F(1,561.64)=101.35, p<.0001;accuracy: χ²(1)=218.44, p<.001). Again, there was a main effect of fluency quintile (RT: F(4,1462.04)=54.27, p<.0001;accuracy: χ²(4)=430.05, p<.001) and a significant interaction between fluency and frequency (RT: F(4,70621.04)=4.61, p=0.001;accuracy: χ²(4)=29.80, p<.001) due to the fact that the slope of the frequency effect decreased slightly in the most fluent readers. Analyses restricted to each fluency quintile showed that, in both RT and accuracy, the frequency effect remained significant.

**Figure 4.**
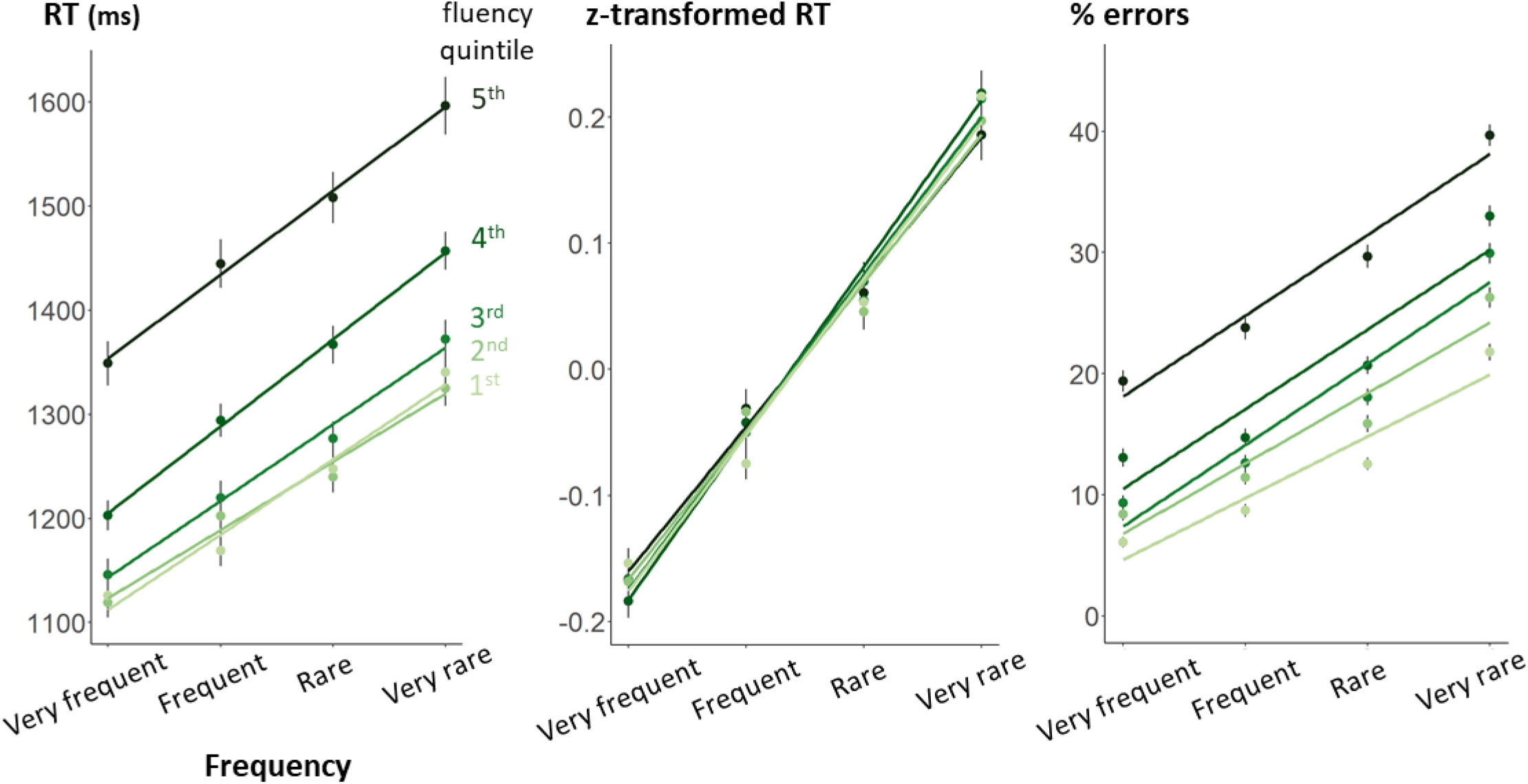
Frequency effects on response times on correct answers (RT), Z-transformed RTs, and error rates. Each point represents the mean RT or error rate as a function of frequency and fluency. Error bars represent the standard error of the mean. The slopes are the linear regression associated with the points.

Again, we used z-transformed RTs to examine whether the reduction in the frequency effect as fluency increased was solely due to faster overall responses (figure 4). The main effect on frequency remained significant, F(1,621.05)=116.36, p<.0001, but the two-way interaction vanished, F(4,72016.56)=1.06, p=0.37. This finding suggests that, once the speed of their responses was taken into account, the frequency effect on RT was actually identical for all students, no matter their fluency level. In other words, frequency appears to be a variable that affects the decision stage and whose amplitude therefore covaries with the standard deviation of RTs, while word length does not.

### Error trials

For words that they know, it may not be surprising that all participants should show a similar word frequency effect. However, as fluency decreased, error rates increased dramatically in the LD task (figures 4, right panel), indicating that less fluent readers also knew fewer words. To understand how unknown words are processed, we performed an analysis of RTs restricted to error trials. This analysis uncovered a significant but reversed frequency effect F(1,747.59)=6.87, p=0.009: on error trials, lower frequency words were responded faster than high-frequency ones, suggesting that, the less frequent a word is, the faster a reader is likely to erroneously classify it as a pseudoword.

### Interaction between length and frequency

Our large sample also allowed us to investigate the interaction between length, frequency, and fluency (figure 5). Our prediction was that these variables should have a 3-way interaction on RTs because (1) fluent readers would show little or no length effect, regardless of frequency, as they consistently rely on the fast lexical route; (2) less fluent readers would show an increasingly marked length effect as word frequency decreases, because lower frequency decreases the probability that they use the lexical route.

**Figure 5.**
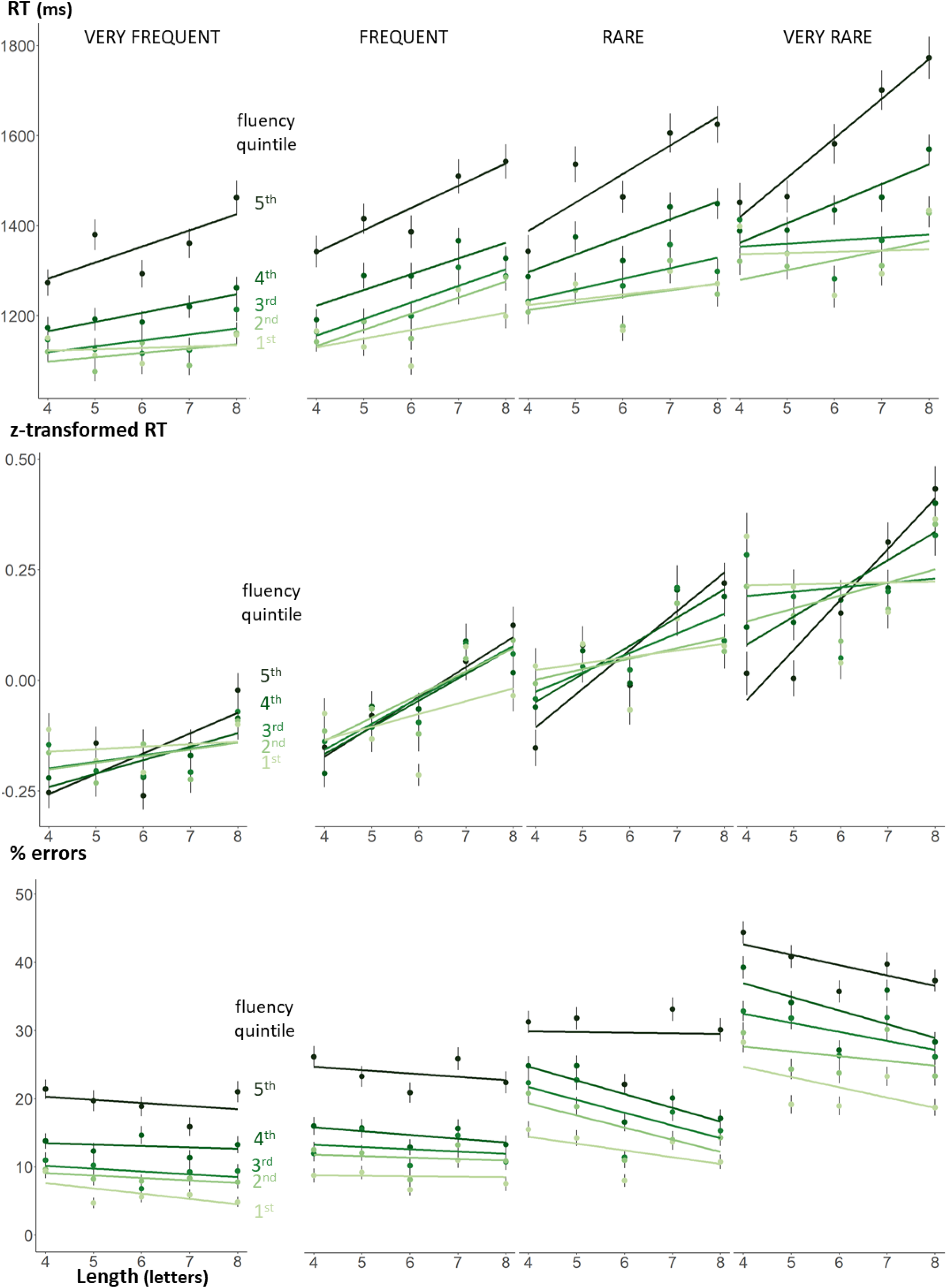
Interaction between length, frequency and fluency on response times on correct answers (RT), Z-transformed RTs and error rates. Each point represents the mean RT or error rate as a function of length, frequency, and fluency. Error bars represent the standard error of the mean. The slopes are the linear regression associated with the points.

In a general linear model restricted to words only, the predicted three-way interaction between fluency quintile, word frequency and length was significant for RT, though not for accuracy (RT: F(4,70602.36)=5.55, p=0.0002 / accuracy: χ²(4)=2.72, p=0.61). Figure 5 shows that RTs were affected by the frequency x length interaction reported earlier, but that, as predicted, this effect decreased as fluency increased. A simple effect analysis on length, separately for each fluency X frequency level, showed a clear trend: the most fluent students exhibited no length effect, no matter the frequency of the words, whereas the least fluent student showed a significant length effect even for very frequent words.

Results on z-transformed RTs were almost exactly the same as for RT, except that, as expected, there was no longer a significant main effect of fluency quintile, F(4,72009.75)=1.30, p=0.27, as well as any significant interaction between word frequency and fluency, F(4,72018.23)=0.99, p=0.41, as previously reported. As the three-way interaction between length, word frequency and fluency level remained significant, our conclusions remained unchanged.

### Processing of pseudowordss

We designed our pseudoword traps for two-by-two comparisons: orthographic traps versus words approximations, transpositions versus double substitutions, and mirror versus single substitutions. Figure 6 shows the data for these comparisons, which we consider in turn.

**Figure 6.**
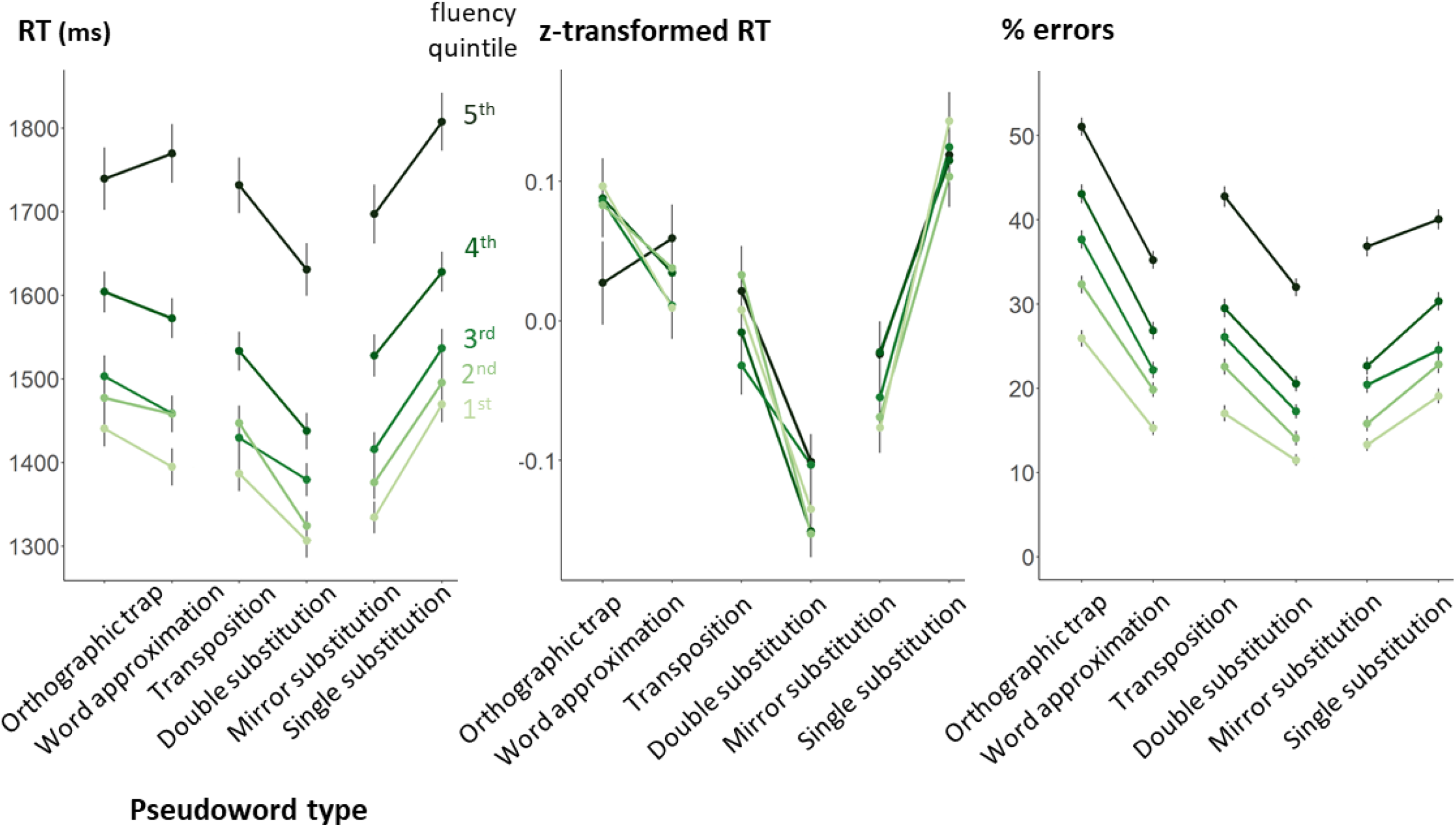
Responses to pseudowords: response times on correct answers (RT), Z-transformedRTs, and error rates

### Impact of orthographic traps

Our mixed effect analysis only exhibit a significant main effect of pseudoword type on accuracy (χ²(1)=32.42, p<.001), not on RT (F(1,147.39)=0.03, p=0.86), meaning that when they were correct, students were equally fast to classify orthographic traps and word approximations. We also found a significant effect of fluency quintile on both RT and accuracy (RT: F(4,1427.42)=36.49, p<.001;accuracy: χ²(4)=394.01, p<.001) and a significant interaction between fluency and pseudoword type only on RT (RT: F1(4, 19155.46)=3.07, p1=0.015;accuracy: χ²(4)=2.98, p=0.56), which was preserved in z-transformed RTs, F(4,20395.02)=2.78, p=0.025. This interaction highlighted that word approximations tended to be classified faster than orthographic traps, except in the least fluent students.

### Effect of letter transpositions

Results of our mixed effect models confirmed these expectations. For both RT and accuracy, we found a main effect of pseudoword type (RT: F(1,175.14)=11.86, p<.001; accuracy: χ²(1)=26.70, p<.001) meaning that transpositions were harder to classify than double substitutions. There was also a main effect of fluency quintile (RT: F(4,1402.42)=46.22, p<.001; accuracy: χ²(4)=403.49, p<.001), but no interaction between fluency and pseudoword type (RT: F(4,21276.14)=2.08, p=0.080; accuracy: χ²(4)=3.46, p=0.484). However, looking at z-scores, we found a significant interaction between type and fluency level, F(4,22626.71)=2.40, p=0.048, confirming that the effect gets higher as fluency level increases.

### Processing mirror substitutions

Surprisingly, our results on this type of pseudoword departed from our prediction that mirror substitutions should be more difficult than single substitutions (figure 6). Our mixed effect model showed a significant main effect of pseudoword type on both RT and accuracy (RT: F(1,174.68)=10.86, p=0.001;accuracy: χ²(1)=13.60, p<.001), but with a difference in favor of mirror substitutions. There was a significant main effect of fluency quintile (RT: F(4,1421.47)=46.40, p<.001;accuracy: χ²(4)=384.37, p<.001) and a significant interaction on accuracy only (RT: F(4,20924.34)=0.26, p=0.90;accuracy: χ²(4)=21.95, p<.001). Within each fluency quintile, the faster and more accurate performance with mirror letters reached significance for all but the least fluent students, χ²(1)=3.21, p=0.073 --the converse of our predictions.

What is the reason for this surprising effect? One possibility is that, due to mirror confusions, mirror pseudowords would quickly access the lexicon (e.g. ‘dalance’ would activate ‘balance’), but that top-down feedback from the lexicon would actually facilitate the detection of the erroneous letter ‘d’ – while this lexical input would not be available for control pseudowords such as ‘falance’. To probe the contribution of the lexical route, we tested for an effect of the frequency of the original word. There was no main effect of frequency (RT: F(1,173.68)=0.21, p=0.65 / accuracy: χ²(1)=0.11, p=0.74) nor, crucially, its interaction with pseudoword type (mirror vs control: RT: F(1,172.59)=0.08, p=0.78;accuracy: χ²(1)=0.67, p=0.41). Thus, the lexical route does not appear to contribute much to the processing of those pseudowords, if at all.

A second possibility is that some mirror-letter substitutions violated the orthographic statistics of French. In French, the letter ‘q’ is very rare and is almost always followed by the letter ‘u’. However, when p was substituted with a q, this graphotactic rule was violated in all but one of our mirror-substituted pseudowords containing a ‘q’. Such a violation could have facilitated the rejection of those pseudowords. To test this idea, we removed all items with a ‘q’ substitution and repeated our mixed effect analysis. The main effect of type vanished for both RT and accuracy (RT: F(1,133.26)=2.17, p=0.14;accuracy: χ²(1)=1.40, p=0.237). There was also no interaction with fluency on RT (F(4,15215.41)=1.88, p=0.11), and a minor one on accuracy: χ²(4)=12.91, p=0.012. This finding suggests that our paradoxical effect on RT (more efficient processing of mirror substitutions) was in fact entirely due to the specifics of the letter ‘q’ – and mirror letters did not pose specific difficulties for our participants.

### Predicting fluency score using Lexical Decision results

Our last goal was to see whether the fluency score of each student could be predicted from his or her LD results. To test this prediction, we extracted five LD parameters for each student: median RT; global error rate; slope of the length effect on RT for all the stimuli; slope of the frequency effect on RT for words; and efficiency score, defined as the ratio of accuracy by mean RT (note that this parameter, like fluency, evaluates the number of correct words per unit of time). The slopes of the length and frequency were calculated for each student by taking their median RT on correct answers for each length (resp. for each frequency category), and measuring the coefficient of a linear model fitted to these data.

The results appear in figure 7. Simple linear regressions indicated that all five parameters were predictive of text reading fluency, with LD global error rate being the most predictive one (r²=0.35). A multiple linear regression showed that all five parameters made significant and independent contributions, with the overall r² reaching 0.38. Thus, we conclude that LD can predict about 40% of the variance in a one-minute text fluency reading test.

**Figure 7.**
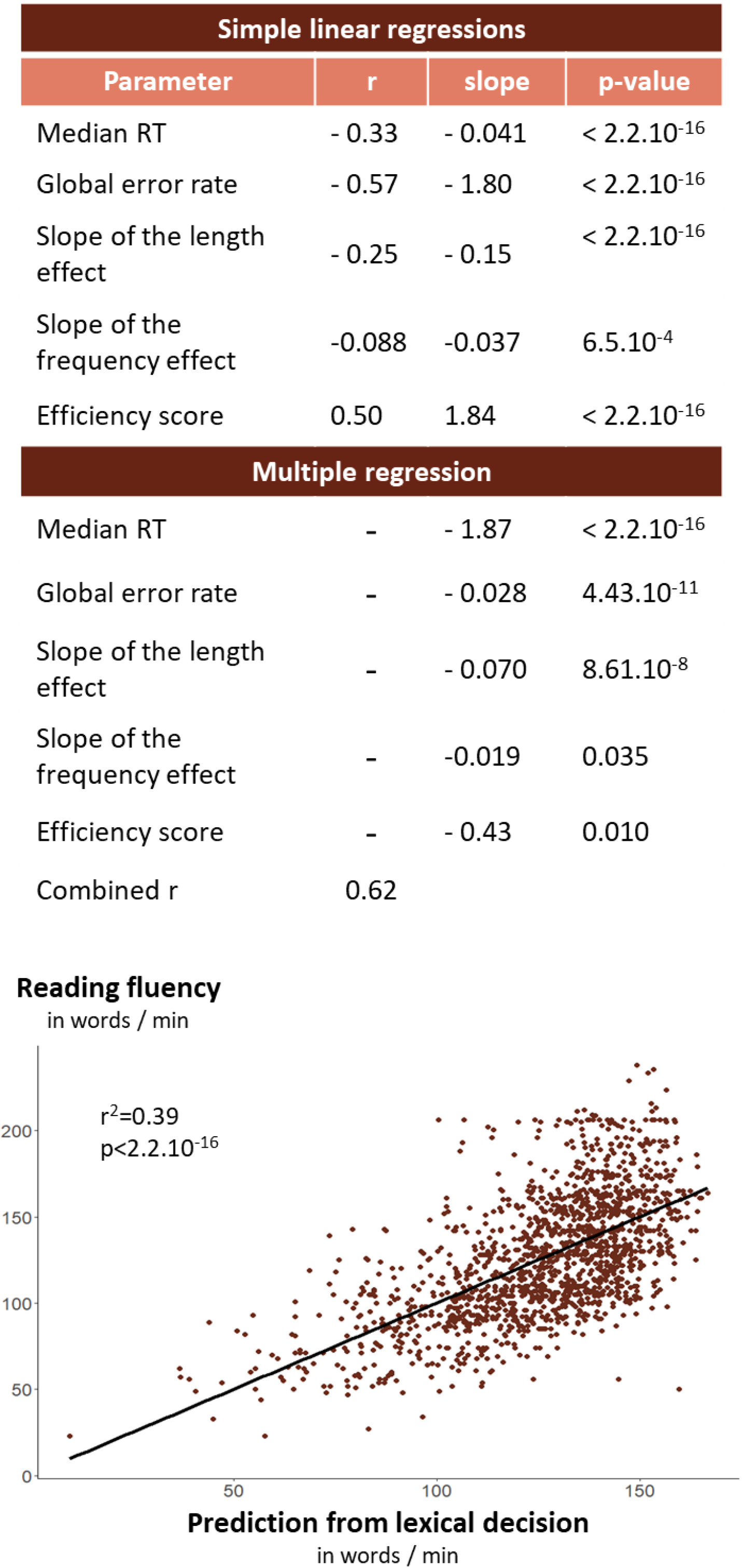
Predicting individual participants’ reading fluency from their results on the lexical decision task.

## Discussion

Our primary goal was to evaluate the lexical decision performance of 6^th^ grade students at different levels of reading fluency, a known predictor of school reading comprehension (Fuchs et al., 2001; Hudson et al., 2005; National Reading Panel, 2000; Pinnell et al., 1995). Our results converges with prior developmental research which compared LD performance across grades, or children with adults, or normal readers with dyslexics – but the originality of the present work is to thoroughly characterized the variability in a large sample of students within a single grade year (6^th^ grade). Our findings corroborate teachers’ reports of a large diversity of reading ability when children arrive in their first year of middle school. Based on student fluency scores, the best performing quintile had already reached adult reading levels, reading more than 200 words per minute, while participants in the worst quintile lagged behind 3^rd^ grade expectations, a striking result with practical implications (Andreu et al., 2022). National statistics suggest that, depending on the definition of literacy, more than 10% of the French populations is struggling with reading, and more than 5% remains functionally illiterate when they leave school and enter active life (Chabanon & Rosenwald, 2020). The present results suggest that the LD test in 6^th^ grade may pick such difficulties at a moment where they might still be corrected.

Our findings are largely compatible with the hypothesis of two different pathways for reading words and pseudowords (Castles, 2006; Coltheart et al., 2001; Di Filippo et al., 2006). In line with previous research, we found a main effect of lexicality, with faster responses to known lexical items. The presence of a significant length effect on pseudoword RTs fits with the hypothesis that pseudowords are deciphered via a slow sublexical route. The decrease in this effect as student’s fluency level increases provides an estimate of the effectiveness of this procedure: the better the students are at reading, the faster their sublexical pathway. On the contrary, when looking at incorrect trials, the length effect on pseudoword RT almost vanished: these pseudowords were misread as words and thus, deciphered through the lexical route.

The impact of length on pseudoword reading contrasts with what was reported by Juphrad and colleagues who found no length effect on pseudoword RT in skilled readers (Juphard et al., 2004a). The type of pseudowords used in the tasks can explain the difference. Juphrad et al’s pseudowords were closest to our trigrams, most of which are orthographically distant from real words, while their words were of very high frequency (minimum 134 per million). Thus, it was far easier for their readers to base their decisions on lexicality, which may explain the absence of a length effect on pseudowords’ RT. In our study, however, the presence of many rare words, as well as the frequent proximity of pseudowords to real words, may have forced participants to rely on decoding.

Concerning words, we found that the length effect on correct RTs was massive for the least fluent students, decreased with fluency, and vanished for most fluent students. This finding is in line with the hypothesis that most fluent students rely on lexical procedures in order to read known words. Conversely, when reading unknown words, i.e. on error trials, all fluency groups exhibit a large length effect, suggesting that they decoded them through their sublexical route. The length effect on accuracy produced a negative trend, which may be due to an interaction with the word neighbors: long stimuli have fewer competitors than short ones, and it is therefore easier to correctly classify them.

Access to the mental lexicon was evaluated by varying the frequency of the word stimuli. As expected, we found a significant frequency effect on both RT and accuracy. The absence of an interaction with fluency in the z-score analysis indicated that, once corrected for greater slowness and variability of responses in the less fluent students, the frequency effect had the same size in all fluency quintiles. This result is in line with those observed by Burani et al (Burani et al., 2002) in younger children, where they showed a frequency effect on naming latencies but no significant interaction between word frequency and grade. It is important to understand that this conclusion applies solely to correct RTs – we also found that less fluent students made many more errors, especially for rare words, suggesting that their lexicon is reduced ; but for words they know, they exhibit a normal frequency effect, suggesting that at this age, all children, regardless of fluency, base their ‘word’ responses in the LD task on an access to the mental lexicon, where frequency is a dominant variable.

Going deeper, we found RT interactions between length and frequency to be dependent on fluency. All readers except the least fluent ones used lexical strategies to read very frequent words. The findings for our best readers regarding lexicality, length and frequency effects were consistent with those described by Araujo et al in Portuguese 3rd to 5th graders (Araújo et al., 2014b). Our poorest reader were similar to previously reported cases of dyslexic readers, as evident by a length effect on RT, thus betraying a strategy of accurate but sublexical reading even for frequent words (Araújo et al., 2014; Zoccolotti et al., 2005). In other words, our poorest readers managed to correctly judge a majority of frequent words, but did so by first identifying them through a slow sublexical reading process.

The third goal of our study was to examine the impact of orthographic distance on pseudoword judgments. In general agreement with previous research, we found that stimuli that were most similar to real French words yielded that highest error rate (Bergmann & Wimmer, 2008; Grainger et al., 2012). Going further, we systematically compared orthographic traps with trigram-based approximations, transpositions with double substitutions, and mirror with single substitutions. We found greater mistakes on orthographic traps than on trigram controls. Such errors are mainly due to regularizations of letter sounds, which is a marker of the use of the sublexical procedure. In line with the literature, we found that transposed letters led to higher RT and error rates than substitutions, and this effect tend to increase between least and most fluent readers (Chambers, 1979; Grainger et al., 2012). It can be seen as a marker of the reliance on the lexical procedure, as all letters are processed in parallel. Consequently, there should be more often confusions in letter position. Finally, looking at mirror and single substitutions, we found a surprising effect: mirror substitutions were faster and more accurately classified than single substitutions. The effect appears to be mainly due to the transformation p→q, When removing these pseudowords, the difference between these two traps also disappears. We conclude that mirror letters, relative to substitutions, do not pose difficulties for French middle-school readers.

Our final goal was to assess the correlation between LD and text reading fluency. We replicate, for adolescents, evidence that accuracy is a better predictor of oral reading ability than RT (Gijsel et al., 2004; van Bon et al., 2004; Yeatman et al., 2021). Previous research has highlighted a correlation of r=0.91 for LD and oral word reading (Yeatman et al., 2021). Here we achieved r=0.62 by using multilinear regression combining median RT, global error rate, slope of the length effect, slope of the frequency effect and efficiency score (thus predicting 38% of the total variance in fluency scores). Our correlation was smaller than the one found by Yeatman et al, perhaps due to a shorter test or to a noisier, more distracting school environment used for measurement. Indeed, the correlation is far from negligible, given that the small number of items seen by each subject, and the fact that the least fluent students produced fewer correct responses on which to collect RTs.

From a practical point of view, these results support the use of LD as a complementary test to oral reading fluency, one that can reveal details of the reading processes in students who show obvious difficulties on the fluency test (Balota et al., 2006; Seidenberg & McClelland, 1989). For example, our poorest readers showed a length effect on RTs to frequent words, suggesting that they know these words but identify them using a sublexical procedure, similarly to dyslexic students (Araújo et al., 2014; Castles, 2006). This insight from LD, which could not be obtained from fluency alone, may help flag students requiring special intervention. To this aim, we have begun to introduce teachers and students with a gamified version of our LD test (figure 8). At the end of this version of the test, students and teachers are provided with a summary of the main results, broken down into two categories: first, the results on words, with examples of errors made in the most frequent category; and second, the results on pseudowords, with examples of the student’s main error type. A tip on how to avoid these types of errors in the future is added at the bottom of the page. In future research, we plan to test the usefulness of this gamified lexical decision test for both students and teachers.

**Figure 8.**
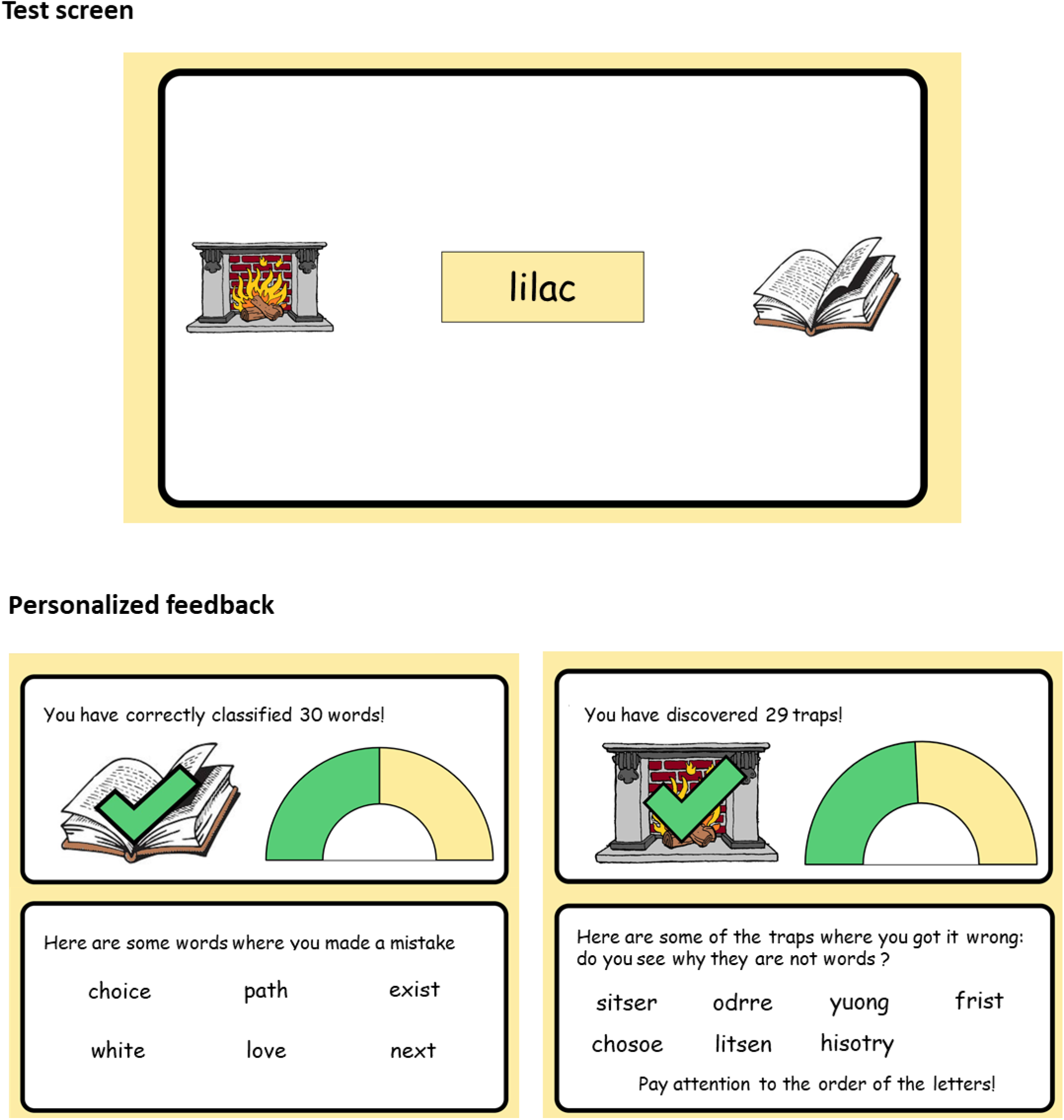
Gamified version of the lexical decision task. Students are asked to send each stimulus Into the dictionary if it is a word, and into the fireplace if it Is a trap. At the end of the session, they receive feedback on their performance, including the number of errors on words and pseudowords and examples of errors. For words, errors are sorted by their frequency. For pseudowords, only the stimuli from the category in which the student made the most errors are displayed. A tip on how to avoid these traps is given.

There are several limitations to this work. As the Ministry of Education carried out the recruitment, we were not able to formally exclude dyslexic students from our participants. Thus, it is likely that part of the group of students with the greatest difficulties was in fact composed of dyslexics. Similarly, we were not able to check for the presence of other learning disabilities that might have disrupted the test, such as attention deficit disorder. Furthermore, the great diversity of items presented to each child left little room for more than 2 or 3 repetitions of the same condition (2 repetitions of each length-type combination for pseudowords, 3 repetitions of each length-frequency combination for words). The high error rates on pseudowords therefore made it impossible to analyze the length effect within each type of pseudoword.

## Conclusion

In summary, our study showed that lexical decision is a useful task to characterize the origins of the large variability in reading abilities in French 6^th^ graders. We suggest that LD cannot replace other oral reading tests such as fluency, but provides an additional assessment that sheds detailed light on the efficiency of the two reading routes, i.e. lexical and sublexical reading procedures. LD has now proven its correlation with other tasks of oral reading. Given it ease of use, LD could prove an excellent tool to differentiate between readers in need of extra practice, and those with more serious deficits marked by an inability to establish an efficient mental lexicon.

## Acknowledgements

We gratefully acknowledge the Direction de l’Evaluation, de la prospective et de la performance (DEPP) of the French National Ministry for Education for its assistance in the deployment of the tests and the data collection and Johannes Ziegler, Mathias Sablé-Meyer and Aakash Agrawal for their advice on data analysis. This work was funded by INSERM, Collège de France, CEA, DEPP, the Collège de France foundation (C.S.) and the Clermont-Tonnerre foundation. M.L. was supported by a CIFRE grant from the CERENE schools and the French national agency ANRT.

## Declaration of interest statement

The author have no conflicts of interest to disclose.

## Ethics statement

This project received ethics approval from the Comite d’Évaluation Éthique d’Établissement (C3E) of the Paris-Saclay University (10/11/2020, project reference: CER-Paris-Saclay-2020-068).

Parents gave their consent for the whole framework of the National Evaluations. Students gave their assent prior to their inclusion in the study.

## Data availability statement

Raw data were collected by the French National Ministry for Education. Participants of this study did not give written consent for their data to be shared publicly, so supporting data is not available.

## Notes

### Competing Interest Statement

The authors have declared no competing interest.

